# EGFR Oncogenes Expressed In Glioblastoma Are Activated As Covalent Dimers And Paradoxically Stimulated By Erlotinib

**DOI:** 10.1101/810721

**Authors:** Matthew O’Connor, Jie Zhang, Sandra Markovic, Darlene Romashko, Andrei Salomatov, Noboru Ishiyama, Roberto Iacone, Alexander Flohr, Theodore Nicolaides, Alexander Mayweg, David Epstein, Elizabeth Buck

## Abstract

Mutation of both the intracellular catalytic domain and the extracellular domain of the receptor for epidermal growth factor (EGFR) can drive oncogenicity. Despite clinical success with targeting EGFR catalytic site mutations, no drugs have proven effective in patients expressing allosteric extracellular domain EGFR mutations, including glioblastomas (GBM) where these mutations are highly expressed. We define the molecular mechanism for oncogenic activation of families of extracellular EGFR mutations and reveal how this mechanism renders current generation small molecule ATP-site inhibitors ineffective. We demonstrate that a group of commonly expressed extracellular domain EGFR mutants expressed in GBM is activated by disulfide-bond mediated covalent dimerization, collectively referred to as locked dimerization (LoDi) EGFR oncogenes. Strikingly, small molecules binding to the active kinase conformation (Type I), but not those binding to the inactive kinase conformation (Type II), potently inhibit catalytic site mutants, but induce covalent dimerization and activate LoDi-EGFR oncogenes, manifesting in paradoxical acceleration of proliferation.

**Significance:** Our data demonstrate how the locked-dimer mechanism of EGFR oncogenesis has a profound impact on the activity of small molecule inhibitors. This provides a mechanistic understanding for the failure of current generation EGFR inhibitors to effectively treat LoDi-EGFR mutants in GBM, and sets guidelines for discovery of selective LoDi-EGFR inhibitors.

## Introduction

Mutations of EGFR that result in oncogenic activity can occur either at the ATP site within the intracellular kinase domain or at allosteric sites within the extracellular domain. Non-small cell lung cancer tumors nearly exclusively express EGFR kinase domain mutations while GBM tumors nearly exclusively express allosteric extracellular domain EGFR mutations (1,2). While oncogenic point mutations that affect the catalytic domain of ErbB receptor tyrosine kinases (RTKs) have been successfully exploited through personalized medicines in oncology (2), there are currently no drugs that have proven clinically effective against the group of allosteric extracellular domain EGFR mutations expressed in GBM.

Mutations of EGFR in GBM nearly exclusively affect the extracellular CR1 and CR2 domains. CR1 and CR2 are cysteine-rich regions characterized by the presence of a series of intramolecular disulfide bonds (3,4). In the absence of ligand EGFR is monomeric and held in an auto-inhibited conformation through interactions that tether the CR1 and CR2 domains (5,6). The binding of ligand to the extracellular domain unleashes the tethered conformation and promotes receptor dimerization through the CR1 and CR2 domains, an event that triggers dimerization and activation of the intracellular catalytic domains (3,4). Therefore, the CR1 and CR2 domains serve to both tether the receptor in an auto-inhibited conformation in the absence of ligand and facilitate interactions at the dimer interface in the presence of ligand.

GBM mutations affecting the CR1/2 domains include the large genomic deletion of exons 2-7 that produces EGFR-Viii. EGFR-Viii is constitutively active in the absence of ligand, exhibits sustained signaling that is resistant to downregulation, and is both transforming and tumorigenic (7–10). EGFR-Viii expression is associated with metastasis and with poor long term overall survival (11). Recent sequencing data has revealed that EGFR-Viii is just one of a group of allosteric mutations affecting the extracellular CR1/2 domains of EGFR in GBM (1). Other mutations include EGFR-Vii and mis-sense mutations at position EGFR-A289, mutants that have also been shown to exhibit oncogenic properties. GBM tumors can co-express multiple oncogenic EGFR allosteric mutations (10).

The expression of multiple EGFR mutants that give rise to transforming and tumorigenic activity makes these receptors especially attractive drug targets for the treatment of GBM. However, drugs that are effective against tumors expressing EGFR catalytic domain mutations have been shown to be ineffective against tumors expressing allosteric EGFR mutations. (12–15). A group of small molecule EGFR inhibitors approved for the treatment of lung cancers expressing EGFR ATP-site mutants (erlotinib, gefitinib, afatinib) have undergone intensive clinical investigations, involving >30 clinical trials and >1500 patients, however all failed to produce clinical benefit, even within a subset of tumors selected for EGFR-Viii expression.

Strikingly, some evidence even suggested that erlotinib promoted disease progression. A phase 2 study evaluating erlotinib in combination with radiation and temozolomide showed median progression free survival (mPFS) and median overall survival (mOS) of 2.8 months and 8.6 months, as compared to 6.9 months and 14.6 months for patients receiving radiation and temozolomide alone (16). Another randomized phase II trial with erlotinib showed that patients who received erlotinib, including those whose tumors expressed EGFR-Viii, performed worse by a number of parameters than those patients who received standard of care therapy (14).

Several recent preclinical studies have modeled the ineffectiveness of erlotinib for inhibiting EGFR-Viii (17,18). This is evidenced in part by observations for poorer binding kinetics for erlotinib to EGFR-Viii compared to either EGFR catalytic site mutants or EGFR-WT. However, there has been no mechanism proposed to explain this effect. Furthermore, there remains a gap in the current understanding of the mechanism responsible for oncogenic activation of other extracellular domain EGFR mutants, and if they would exhibit similar receptor pharmacology. We therefore, sought to understand the mechanism of receptor activation and its impact on EGFR inhibitor activity. Our novel findings demonstrate that covalent homodimerization, driven through disulfide bond formation at the extracellular dimer interface, is a unifying mechanism of activation for a family of allosteric mutant EGFR oncogenes. Importantly, this mechanism reveals an allosteric alteration of the pharmacology profile for small molecules binding to the intracellular ATP-site of these oncogenic kinases that has profound consequences on the activity for current generation small molecule EGFR inhibitors.

## Results

### Oncogenic mutations of EGFR affecting allosteric sites in the extracellular domain are commonly expressed in glioblastoma

RNA sequencing of EGFR in GBM tumors has revealed a spectrum of alterations (1). Commonly expressed alterations include large exonic deletions affecting the extracellular domain such as EGFR-Viii and EGFR-Vii. EGFR-Viii results from deletion of exons 2-7, and is expressed by nearly 20% of GBM tumors, Figure 1A/B. EGFR-Vii results from deletion of exons 14-15, and is expressed by 3% of glioblastoma tumors. Another alteration results in deletion of exons 12-13, which we herein define as EGFR-Vvi, and is expressed by greater than 30% of GBM tumors, Figure 1A/B. Genomic missense mutations affecting EGFR are also commonly expressed in glioblastomas, observed in 48% of cases. However, in contrast to lung adenocarcinoma tumors, where missense mutations and short indels nearly exclusively affect the intracellular kinase domain, missense mutations expressed in GBM nearly exclusively affect allosteric sites in the extracellular domain. Mutation of A289 in the CR1 region of the extracellular domain is the most commonly occurring missense mutation in GBM, expressed by nearly 12% of tumors. Interestingly, many GBM tumors express multiple extracellular domain EGFR mutations, Figure 1A. Among 31 tumors that express EGFR-Viii, 18 co-express Vvi, and only 4/31 tumors express EGFR-Viii in the absence of EGFR-Vvi, EGFR-Vii, or an EGFR short variant mis-sense mutation of the extracellular domain.

**Figure 1.**
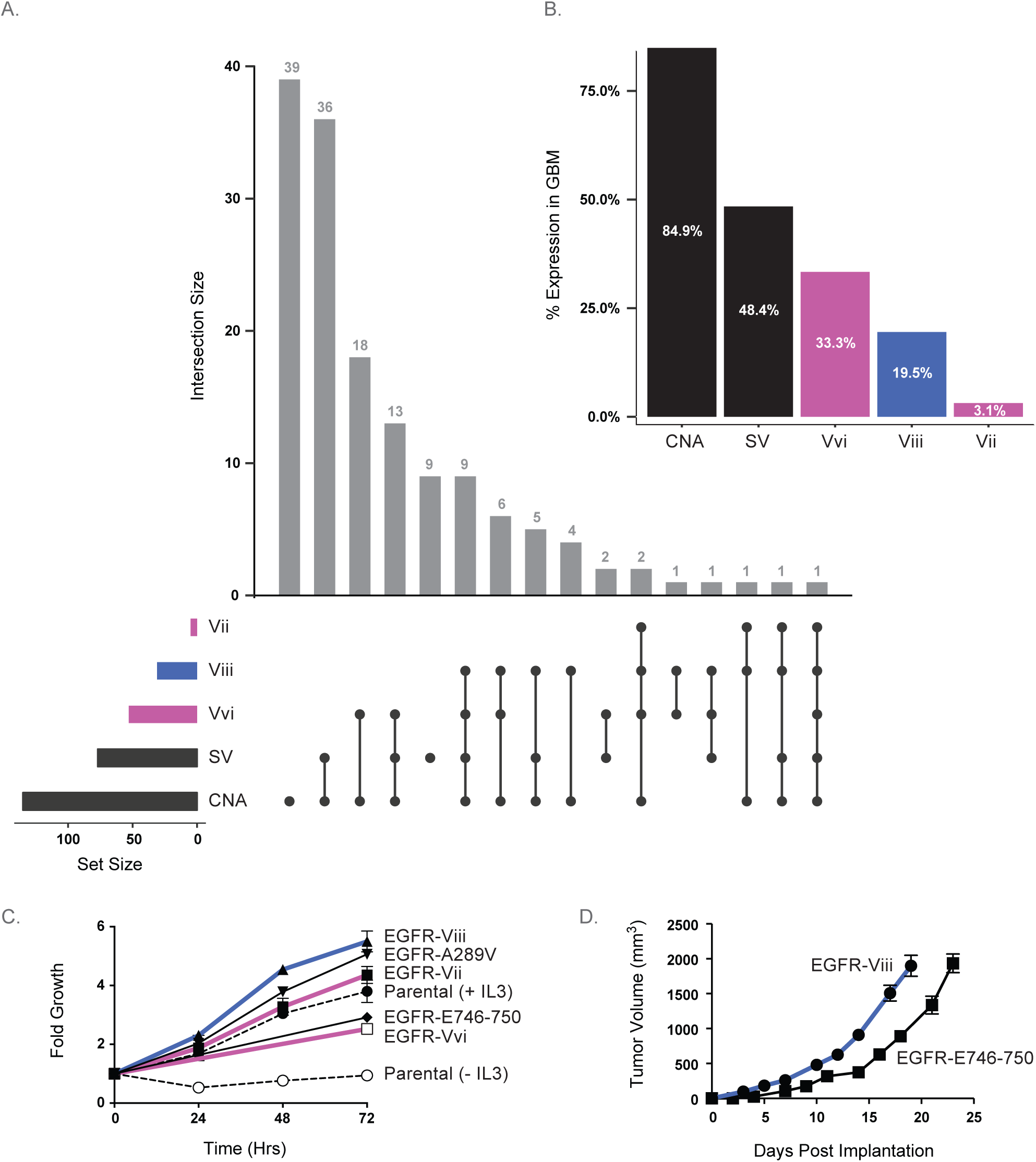
Glioblastoma tumors commonly express EGFR oncogenic mutations affecting the extracellular domain. **A.** UpSetR (35) visualization of co-expression of short variants (SV), large exonic deletions Vii, Viii, Vvi, and regional gain or focal amplification events (CNA) in glioblastoma cases from Brennan et al (1). Mutations that were expressed with an allelic frequency of greater than 1% were considered positive. **B.** Genomic alterations in glioblastoma cases from Brennan et al (1), prevalence. **C.** Proliferation of parental BaF3 cells cultured in the presence or absence of IL-3 or proliferation of BaF3 cells transformed with EGFR-Viii, EGFR-Vii, EGFR-Vvi, EGFR-A289V or EGFR-E746-750, and cultured in the absence of IL-3. Proliferation was measured over a 72-hour period. **D.** Growth of tumors in athymic nude mice with subcutaneous implantation of BaF3 cells transformed with EGFR-E746-750 or EGFR-Viii.

Previous studies have demonstrated that extracellular domain EGFR mutations in GBM, including EGFR-Viii, EGFR-Vii, and EGFR-A289V are oncogenic, (10,19,20). To confirm the oncogenic properties for this family of EGFR mutations, we assessed their ability to transform BaF3 cells to IL3 independence in comparison to the clinically validated oncogenic EGFR ATP-site mutation EGFR-E746-750 that is expressed in lung adenocarcinoma. Here we confirm the transforming properties for EGFR-Viii, EGFR-Vii, and EGFR-A289V and demonstrate that EGFR-Vvi, the most commonly expressed mutation in GBM, also transforms BaF3 cells to proliferate in the absence of IL3, Figure 1C. This family of extracellular domain mutants transforms BaF3 cells to proliferate to a similar or greater extent versus EGFR-E746-750. Both EGFR extracellular domain and ATP-site mutants also promote tumor growth in vivo, Figure 1D. EGFR-Viii and EGFR-E746-750 BaF3 transformants readily form tumors in mice, with somewhat shorter latency for EGFR-Viii versus EGFR-E746-750 allografts. These data support the oncogenicity for the family of extracellular domain EGFR mutations in GBM, to a similar extent as EGFR ATP-site mutations in lung cancer.

### The oncogenicity of allosteric EGFR oncogenes in GBM is mediated by disulfide-linked covalent homodimerization

Two cysteine rich regions in the extracellular domain, CR1 and CR2, mediate the auto-inhibitory intramolecular interface for an inactive EGFR monomer. These same cysteine rich regions also mediate the intermolecular interface for a ligand-activated EGFR dimer (3,4), as shown in Figure 2A and 2B. A common feature of EGFR mutations expressed in GBM is their location in the CR1 or CR2 cysteine rich regions, Figure 2C. Mapping the EGFR-Viii and EGFR-Vii mutations onto a protein structure revealed that auto-inhibitory tethering interactions would be disrupted, Figure 2A. Tethering is mediated primarily by a highly conserved triangular salt bridge and hydrogen bond network formed by Y270 (CR1), D587 (CR2), and K609 (CR2) (EGFR numbering), Figure 2A. In EGFR-Viii Tyr270 of the YDK triad is lost, while for EGFR-Vii both Asp587 and Lys609 are lost, thereby preventing adoption of the inactive tethered position and promoting formation of an extended conformation that is poised for dimerization. Indeed, this mechanism has been described for several EGFR mutations expressed in GBM (21). Previous studies have also shown that the oncogenic activation of EGFR-Viii may be attributed to disulfide bond-mediated covalent dimerization of the extracellular domain (22,23). Mapping cysteine bonding in the CR1 and CR2 domains of the EGFR-Viii, -Vii, and - Vvi truncations reveals that each truncation will disrupt one or more intramolecular disulfide bonds, resulting in the presentation of free cysteines at the extracellular dimer interface and the potential for intermolecular disulfide bond formation between receptors upon dimerization, Figure 2B/C. In EGFR-Vii, mutation results in disruption of three intramolecular disulfide bonds in the CR2 domain, Cys539-Cys538, Cys620-Cys628 and Cys624-Cys636 bonds, leaving Cys538, Cys628 and Cys636 as free cysteines. Loss of exons 14-15 in EGFR-Vvi disrupts the Cys539-Cys538 bond, leaving Cys538 as a free cysteine. All free cysteines are situated at the EGFR dimer interface, Figure 2B. Inspection of the site of EGFR-A289V mutation within the x-ray structure of the EGFR ectodomain reveals it is also situated in close proximity to intramolecular disulfide bonds at the dimer interface, Figure 2B. We speculated that this mutation might disrupt neighboring disulfides, forming free cysteines at the dimer interface. Indeed, mutations that are adjacent to a disulfide bond in the third Ig-like domain of FGFR2 have been shown to disrupt this bond, forming free cysteines and conferring a covalently dimerized and activated receptor (24). EGFR-A289 is less than 10 angstroms from the Cys-264-Cys291 bond, and alterations at this site might prevent formation of this disulfide, resulting in presentation of free cysteines at the CR1 dimer interface region.

**Figure 2.**
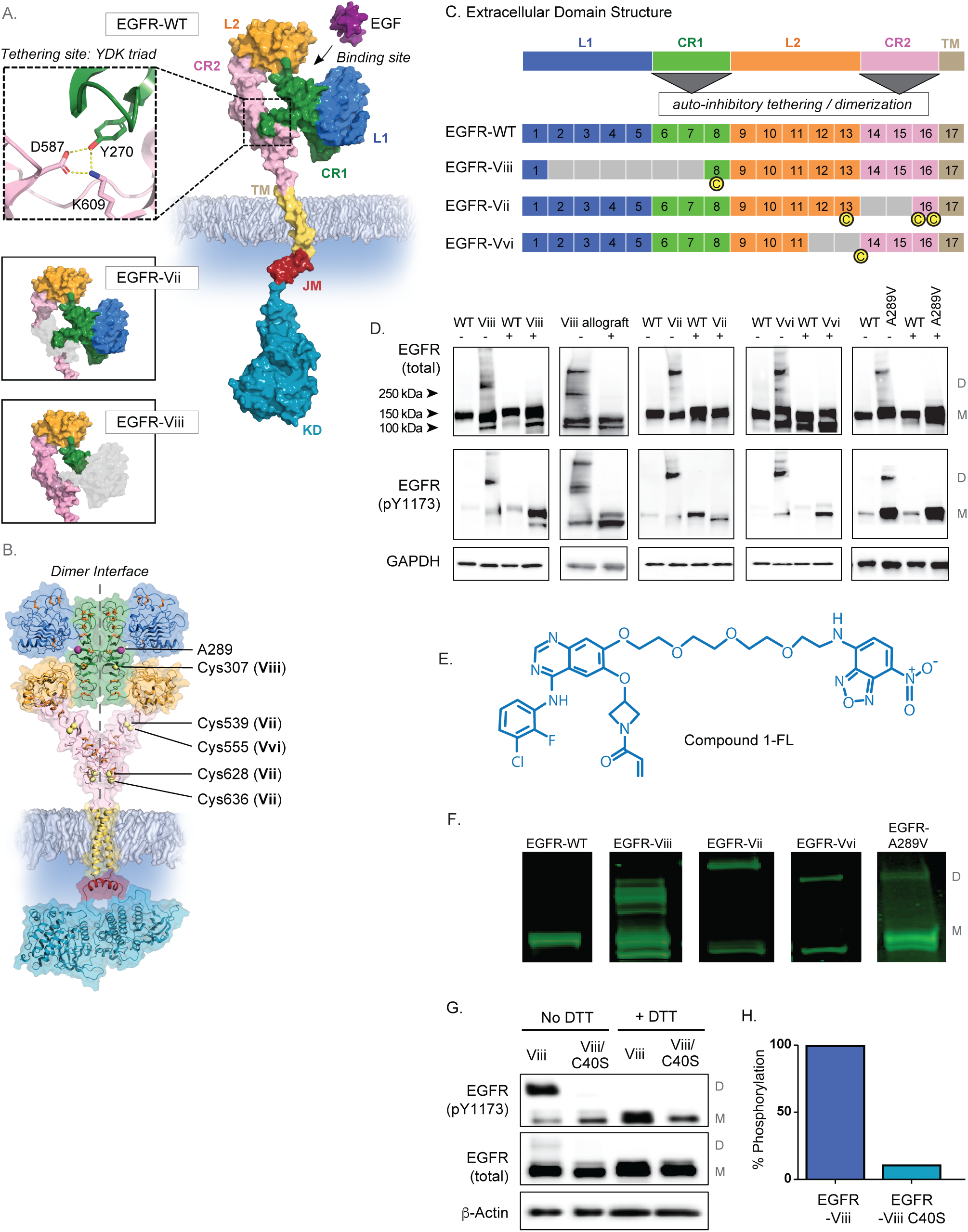
Mutations affecting the extracellular domain of EGFR occur at sites functioning in auto-inhibition and dimerization and result in covalent activation. **A.** Molecular surface representation of EGFR describing the extracellular domain in the tethered conformation (PDB ID: 4UV7) and resolved using PyMOL (http://www.pymol.org/). The L1 domain is shown in blue, CR1 in green, L2 in orange, and CR2 in pink. Regions truncated in the EGFR-Viii, -Vii, and - Vvi mutants are shown in grey. Within the model for EGFR-WT a detailed schematic for the YDK triangular hydrogen bonding network forming a key tethering interaction is provided. TM, JM and KD indicate transmembrane, juxtamembrane and kinase domains, respectively. **B.** Ribbon model of EGFR in the dimerized conformation based on representative atomic coordinates (PDB ID: 3NJP, 2KS1 and 4G5P). The dimer interface is noted. Intramolecular disulfide bonds are shown as orange sticks. The positions of free cysteines generated by the EGFR-Viii, EGFR-Vii and EGFR-Vvi mutants are noted as yellow spheres. The position of the A289V mutation is indicated. **C.** Schematic representation of the extracellular domain structure of EGFR. Exons coding the L1, L2, CR1, and CR2 regions are noted, as are regions functioning in auto-inhibition and dimerization. Exons truncated in the EGFR-Viii, EGFR-Vii, and EGFR-Vvi mutants are shown in grey, and positions of free cysteines resulting from these mutations are noted by C in yellow below each schematic. **D.** The expression of total and phosphorylated forms of LoDi-EGFR mutants EGFR-Viii, EGFR-Vii, EGFR-Vvi, and EGFR-A289V versus EGFR-WT. Expression was detected by resolving proteins by SDS PAGE in the presence (+) or absence (-) of reductant (DTT) followed by western blotting using antibodies to recognize total and phosphorylated EGFR. Lysates for mutants are derived from Ba/F3 transformants, and lysates for EGFR-WT are derived from A431 cells. **E.** Molecular structure of the Compound 1-FL probe. **F.** Fluorescence imaging of proteins derived from EGFR-WT (A431) or EGFR mutant (BaF3 transformants) cells and resolved by SDS PAGE following treatment with Compound1-FL probe for 20 minutes. **G.** The expression of total and phosphorylated forms of EGFR-Viii and EGFR-Viii/C40S mutation. Expression was detected by resolving proteins by SDS PAGE in the presence or absence of reductant (DTT) followed by western blotting using antibodies to recognize total and phosphorylated EGFR. Protein lysates were derived from U87MG cells expressing each of the EGFR mutants. **H.** Expression level of phosphorylated EGFR dimer in EGFR-Viii/C40S mutant as a percentage of phosphorylated dimer in the EGFR-Viii mutant, determined by quantitative densitometry.

To test whether free cysteines positioned at the dimer interface for allosteric EGFR mutants could promote covalent dimerization and activation we performed western blotting for EGFR under non-reducing conditions to allow the visualization of covalently linked dimers. In A431 tumor cells that express high levels of EGFR-WT, EGFR is resolved as a single band migrating near the 150KDa marker even under non-reducing conditions, consistent with monomeric WT-EGFR, Figure 2D. In contrast, we found evidence for covalent dimerization for each of the extracellular domain EGFR mutant oncogenes. For EGFR-Viii and other mutants we resolved two bands, one migrating near the 150kDa marker, consistent with monomer, and a second band with a higher molecular weight migrating above the 250kDa marker, consistent with a dimer. We also observed the presence of a minor band at a much higher molecular weight, which could represent a multimeric state. Both the higher molecular weight band and the band corresponding to EGFR dimer were reduced to a molecular weight corresponding to monomer when lysates were resolved in the presence of DTT reductant. For EGFR-Viii, we confirmed the identity of monomer and covalent dimer bands along with the multimer band by mass spectrometry (data not shown). For each mutant, we found that while the covalent dimer represented a minor fraction of total receptor levels, nearly all phosphorylated receptor, indicative of the active form, was present as a covalent dimer. We validated the covalent mechanism for EGFR-Viii activation in vivo, Figure 2D. Characterization of tumor lysates derived from EGFR-Viii BaF3 allograft tumors revealed a band corresponding to receptor dimer, along with the higher molecular weight multimer, which could be reduced to the monomeric form in the presence of reductant. Due to the ability of this group of extracellular domain mutants to form locked covalent dimers, we herein refer to this family as Locked-Dimer (LoDi) EGFR oncogenes.

To further validate covalent dimer activation for LoDi-EGFR oncogenes, we developed an active site fluorescent probe that binds selectively and covalently to the active site cysteine of EGFR, Compound1-FL, Figure 2E. Compound1-FL was designed based upon a previously described active site fluorescent probe (18). When A431 cells expressing WT-EGFR were treated with Compound1-FL and lysates were subjected to SDS-PAGE in the absence of reductant, we observed fluorescence from a single band corresponding to the molecular weight of monomeric EGFR-WT, Figure 2F. In contrast, when BaF3 cells expressing each of the LoDi-EGFR mutants were treated with Compound1-FL, we observed fluorescence from two bands, one corresponding to the molecular weight of the monomeric mutant and the other corresponding to the molecular weight for the covalently dimerized mutant.

Covalent dimerization and constitutive activation of LoDi-EGFR mutants is dependent on the free cysteine generated in the extracellular domain. When expressed in cis with EGFR-Viii, mutation of the free cysteine Cys307 (Cys40 using EGFR-Viii numbering) to Ser blocks the ability of EGFR-Viii to covalently dimerize, Figure 2G. The Cys307Ser (C40S) mutation results in attenuation of EGFR-Viii phosphorylation by >90% (Figure 2H), demonstrating how the free cysteine in the extracellular domain is critically important for both covalent dimerization and oncogenic activation.

### Covalently activated LoDi-EGFR oncogenes exhibit altered ATP-site pharmacology, and are paradoxically activated by current generation EGFR inhibitors including erlotinib

The primary amino acid sequence of the catalytic domain is unaffected by oncogenic LoDi-EGFR mutations within the extracellular domain. We sought to test whether covalent dimerization of LoDi-EGFR mutants might globally affect the function of the catalytic domain and the pharmacology profile for small molecules that bind to the intracellular ATP-site. We utilized potent, ATP competitive tyrosine kinase inhibitors (TKIs) as sensitive probes of EGFR kinase domain structure and function. For our initial analysis, we selected the potent EGFR inhibitor, erlotinib, for evaluation. Surprisingly, we found that erlotinib enhanced the formation of covalent dimers for all LoDi-EGFR mutants, Figure 3A. These effects were dose-dependent and observed at concentrations as low as 1nM, Figure 3B.

**Figure 3.**
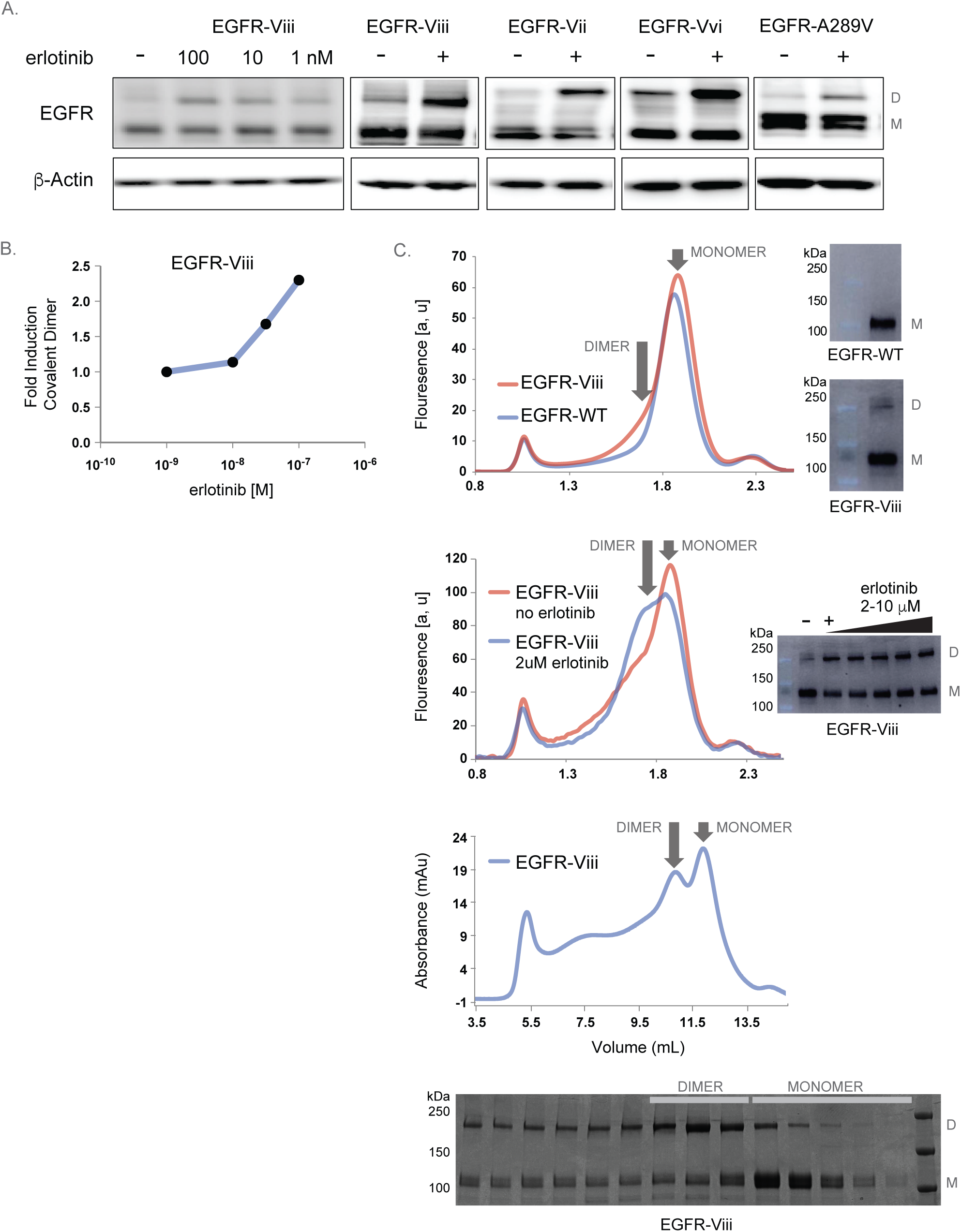
Erlotinib enhances covalent dimerization of LoDi-EGFR oncogenes. **A.** Effect of varying concentrations of erlotinib (EGFR-Viii) or 100nM erlotinib on the levels of monomeric and dimeric -LoDi-EGFR oncogenes. Lysates are derived from U87MG cells expressing EGFR-Viii, EGFR-Vii, EGFR-Vvi, or EGFR-A289V **B.** Quantitation of band density for covalent EGFR-Viii dimer following treatment with varying concentrations of erlotinib. **C.** *Upper panel*: Fluorescence-detection size-exclusion chromatograms of detergent-solubilized cell lysate of HEK293F cells expressing EGFR-WT GFP fusion (red) and EGFR-Viii GFP fusion (blue). While the FSEC profile of EGFR-WT GFP fusion suggests a monodisperse monomeric protein, the profile of EGFR-Viii GFP fusion suggests some presence of a dimer labelled with the arrow. The insets show in-gel fluorescence of EGFR-WT GFP fusion protein (upper) and EGFR-Viii GFP fusion protein (lower). SDS-PAGE was performed under non-reducing conditions. *Middle panel*: Fluorescence-detection size-exclusion chromatograms of EGFR-Viii GFP fusion protein expressed with (blue) and without erlotinib (red) in the culture medium. The inset shows in-gel fluorescence of detergent solubilised cell lysates of HEK293F cells treated with increasing erlotinib concentration in expression medium. SDS-PAGE was performed under non-reducing conditions. *Lower panel:* Size-exclusion chromatogram of EGFR-Viii expressed in the presence of erlotinib (upper panel). Detergent-solubilised and purified EGFR-Viii was subjected to size-exclusion chromatography on a Superose 6 10/300 Increase column equilibrated in buffer containing 0.2 mM dodecylmaltoside. SDS-PAGE analysis of eluted fractions, performed under non-reducing conditions (lower panel). The arrows mark positions of EGFR-Viii dimer and monomer.

To further validate the propensity for erlotinib to induce covalent dimerization, we expressed EGFR-WT and EGFR-Viii C-terminally fused to GFP in HEK293F cells. Detergent solubilized cell lysates containing expressed EGFR variants were analyzed by fluorescence-detection size inclusion chromatography (FSEC) and by in-gel fluorescence. EGFR-WT GFP-fusion resolved as a monomer when assessed using in-gel fluorescence, and eluted as a single monomeric peak in FSEC, Figure 3C (top panel). However, consistent with our measurements of covalent dimerization for EGFR-Viii by both western blotting and by the active site fluorescent probe, EGFR-Viii GFP-fusion resolved as both a dimer and monomer band when assessed using in-gel fluorescence and exhibited a shoulder that would correspond to the dimer when assessed through FSEC. Treatment of EGFR-Viii GFP-fusion with erlotinib resulted in increased presence of the dimer on the gel, Figure 3C (middle panel). Further, there was a shift from a monomeric to a dimeric peak observed by fluorescence chromatography when GFP-fused EGFR-Viii was treated with erlotinib. EGFR-Viii dimer was also detected when the protein was expressed in the presence of erlotinib on a larger scale and subsequently extracted from the membrane using detergent and purified. Size exclusion chromatography of erlotinib-treated EGFR-Viii clearly distinguished a dimer and monomeric elution peak, which was confirmed when fractions were analyzed by gel electrophoresis, Figure 3C (bottom panel).

The propensity for erlotinib to promote covalent dimerization of LoDi-EGFR mutants is associated with enhanced functional activation of LoDi-EGFR. Treatment with sub-saturating concentrations of erlotinib induced phosphorylation of the LoDi-EGFR mutants EGFR-Vii, EGFR-Viii, and EGFR-A289V, Figure 4A. 10nM erlotinib induced the phosphorylation of LoDi-EGFR in cells expressing both LoDi-EGFR-Viii and LoDi-EGFR-Vii, while for EGFR-Vii, inhibition of phosphorylation was observed at 1uM, a very high concentration of erlotinib (Figure 4A). Further, when cells expressing either LoDi-EGFR-Viii, LoDi-EGFR-Vii, or LoDi-EGFR-Vvi were treated with erlotinib, we found that all of these cells showed enhanced phosphorylation compared to untreated control cells (Figure 4B).

**Figure 4.**
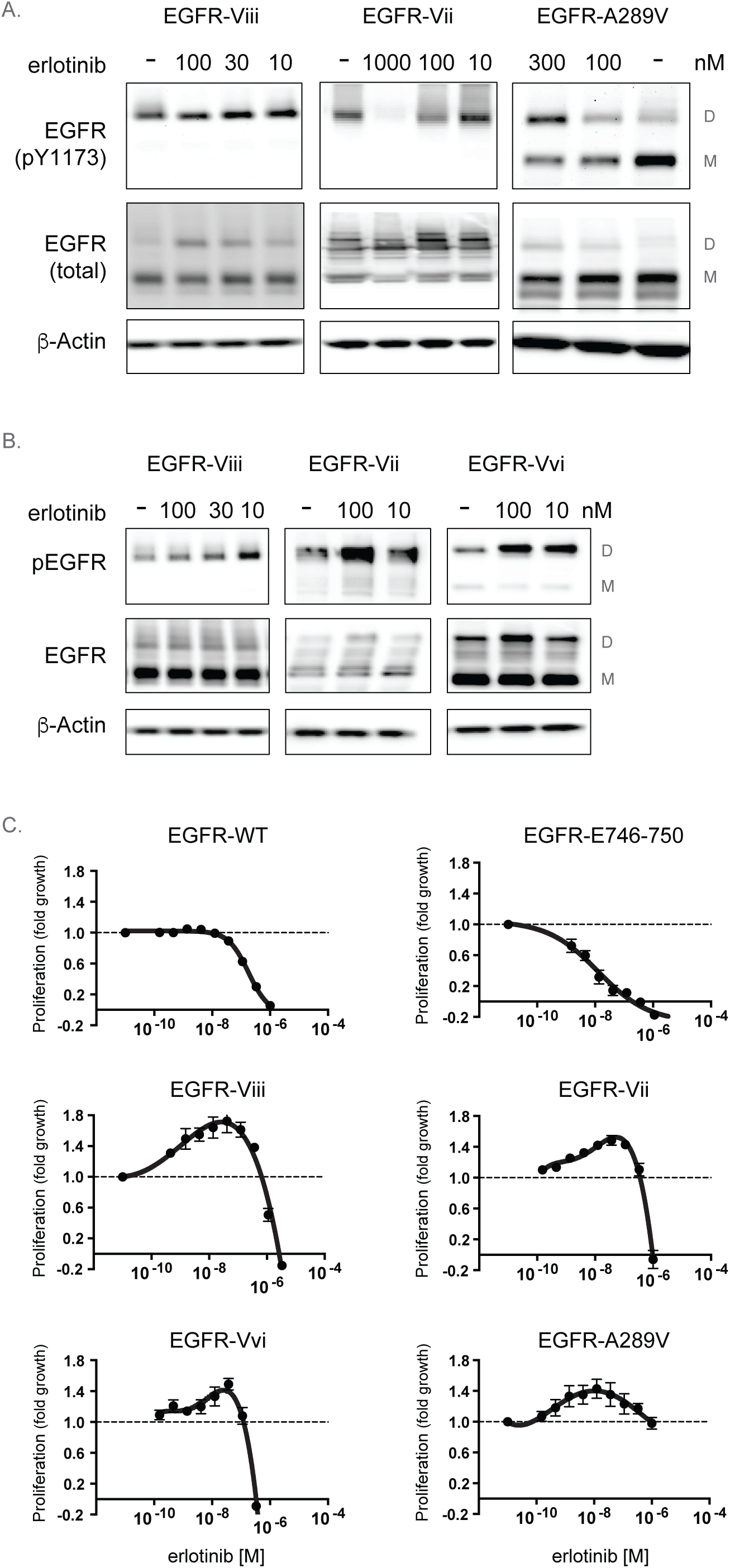
Sub-saturating concentrations of erlotinib stimulate the phosphorylation of covalently dimerized receptors in cells expressing EGFR-Viii, EGFR-Vii, and EGFR-A289V. **A.** Effect of varying concentrations of erlotinib on monomeric and dimeric levels of total and phosphorylated EGFR-Viii (left panel), EGFR-Vii (middle panel), and EGFR-A289V (right panel). Proteins were derived from U87MG cells expressing each of the EGFR mutants and resolved under non-reducing conditions. **B.** Effect of varying concentrations of erlotinib treatment, followed by 30-minute washout, on total and phosphorylated EGFR levels in cells expressing EGFR-Vii or EGFR-Vvi. Proteins were derived from U87MG cells expressing each of the EGFR mutants and resolved under non-reducing conditions. **C.** Effect of varying concentrations of erlotinib on the proliferation of A431 cells (EGFR-WT) or BaF3 cells transformed with the EGFR mutants EGFR-E746-750, EGFR-Viii, EGFR-Vii, EGFR-Vvi, and EGFR-A289V.

In contrast to the potent anti-proliferative activity of erlotinib against WT-EGFR expressing A431 cells (IC50 = 144nM) or BaF3 transformants expressing the catalytic domain mutant EGFR-E746-750 (IC50 = 12nM), BaF3 transformants expressing LoDi-EGFR oncogenes demonstrate insensitivity to the anti-proliferative effects of erlotinib, Figure 4C. Consistent with the enhanced functional activation of the receptor, erlotinib paradoxically stimulates the proliferation of each LoDi-EGFR oncogene transformant at sub-saturating doses, achieving inhibition only at very high saturating drug levels, Figure 4C.

### The conformation of the ATP-site is linked to the extracellular domain, and to signaling that affects pharmacology for small molecule inhibitors against LoDi-EGFR oncogenes

In order to understand if dimer induction and paradoxical activation is a property of all small molecule EGFR inhibitors, we tested molecules representing different scaffolds, distinct binding modes (e.g., Type I or Type II), and both reversible and covalent binding mechanisms (25) (Figure 5A). We found that small molecules representing both quinazoline and pyrimidine cores (e.g., six quinazolines and five-pyrimidines, respectively) and with both reversible and covalent binding mechanisms (e.g., one-reversible and ten-covalent) could induce covalent dimerization (Figure 5B, and supplementary data).

**Figure 5.**
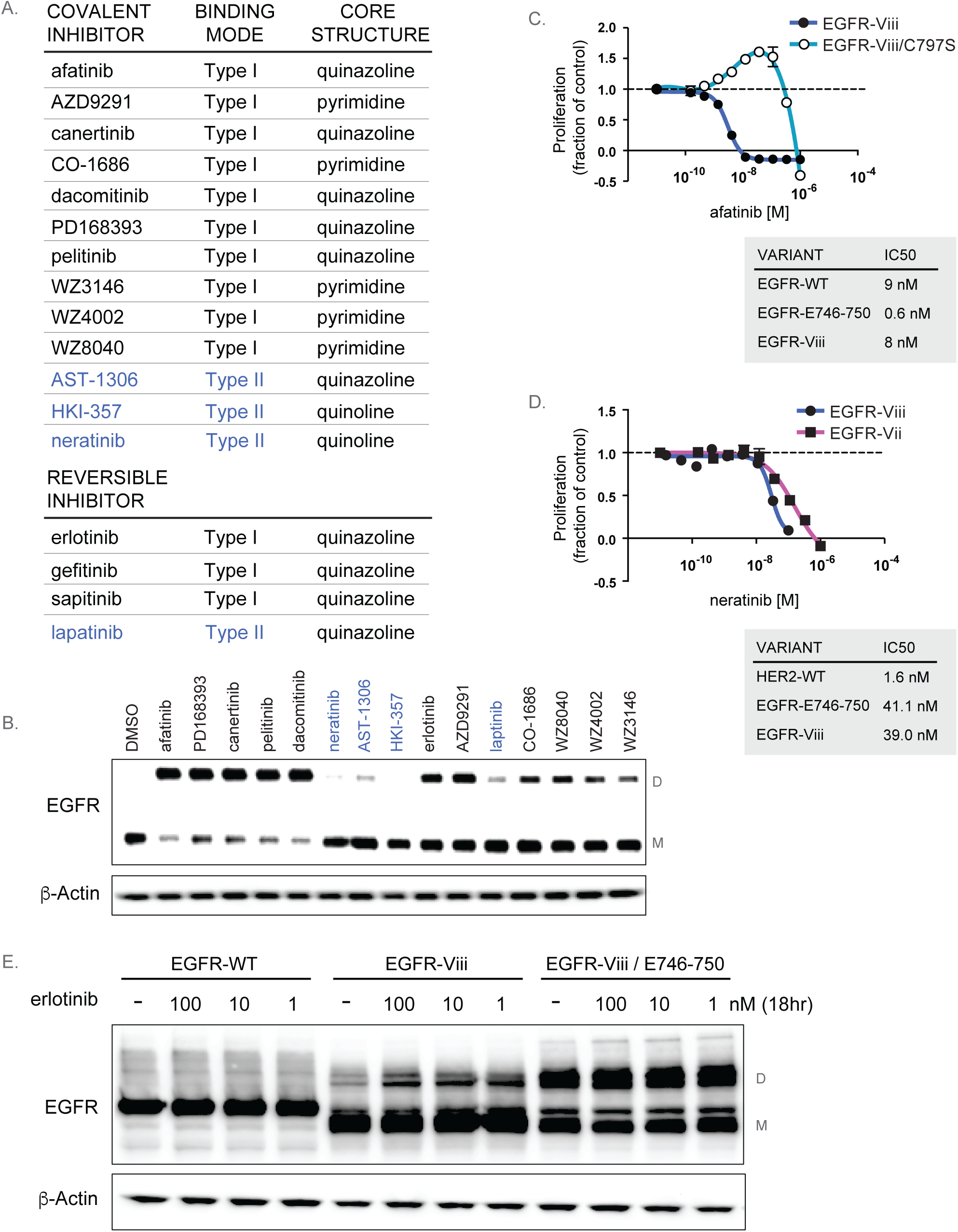
Small molecules or mutations that stabilize the active conformation of EGFR promote covalent dimerization and paradoxical activation of LoDi-EGFR oncogenes. **A.** A selected panel of small molecule ErbB inhibitors described according to binding mode, reversibility or covalency, and core structure. Effect of a panel of diverse EGFR inhibitors on levels of monomeric or covalently dimerized EGFR in U87MG cells expressing EGFR-Vii. Proteins were resolved in the absence of reducing agent to capture covalent dimers. **B.** Effect of varying concentrations of afatinib on the proliferation of BaF3 cells transformed with EGFR-Viii or EGFR- Viii/C797S. Table shown represents cellular antiproliferative IC50 values (nM) of afatinib for EGFR-WT cells (A431), EGFR-E746-750 mutant cells (BaF3 transformants), and EGFR-Viii mutant cells (BaF3 transformants). **C.** Effect of varying concentrations of neratinib on the proliferation of BaF3 cells transformed with EGFR-Viii or EGFR-Vii. Table shown represents antiproliferative cellular IC50 values (nM) of neratinib for HER2-WT cells (BT474), EGFR-E746-750 mutant cells (BaF3 transformants), and EGFR-Viii mutant cells (BaF3 transformants). **E.** Levels of total EGFR for EGFR-WT cells (A431), U87MG cells expressing the EGFR-Viii mutation, or cells expressing the EGFR-Viii mutation in cis with EGFR-E746-750. Effects were measured in the presence of varying concentrations of erlotinib, and lysates were resolved under non-reducing conditions to capture covalent dimers.

Two classes of inhibitors bind to the kinase active site and stabilize either the active conformation (Type I inhibitor binding mode) or the inactive conformation (Type II inhibitor binding mode). Erlotinib, gefitinib, osimertinib, and afatinib exemplify molecules with a Type I binding mode, while lapatinib and neratinib exemplify molecules with a Type II binding mode (25,26). In contrast to the activities observed for the Type I EGFR kinase inhibitors on LoDi-EGFR, EGFR-directed Type II inhibitors do not induce covalent dimerization (Figure 5B, and Supplementary Figure 1). We tested whether other Type I inhibitors that potentiate covalent dimerization of LoDi-EGFR mutants would also show evidence for paradoxical stimulation of proliferation. While the reversible Type I inhibitor gefitinib showed a similar bell-shaped effect on the proliferation of EGFR-Viii BaF3 transformants (Supplementary Figure 2), the covalent Type I inhibitor afatinib exhibited a sigmoidal dose response curve with no evidence for paradoxical stimulation of cell proliferation (Figure 5C). Here the potency for afatinib against EGFR-Viii cells was similar to that observed for A431 EGFR-WT cells (8nM versus 9nM), although this was still >10-fold less potent versus cells expressing the EGFR-E746-750 (IC50 of 0.6nM), the mutation associated with afatinib activity in patients. We suspected this difference in behavior for afatinib versus erlotinib and gefitinib was related to afatinib’s ability to form a covalent interaction with the active site cysteine, C797, wherein this drug would remain bound to the receptor to maintain inhibition of receptor activity even while it promotes covalent dimerization. To test this, we mutated C797 to serine to convert afatinib’s binding mode from covalent to reversible. The C797S mutation, when expressed in cis with EGFR-Viii, translated to a change in afatinib’s phenotype from inhibitor to bell-shaped activator (Figure 5C).

Type II inhibitors, including neratinib, did not potentiate covalent dimerization for LoDi-EGFR mutants. Consistent with this behavior, we found that neratinib inhibited the proliferation of tumor cells driven by LoDi-EGFR oncogenes without any evidence for paradoxical activation (Figure 5D). The anti-proliferative IC50 for neratinib against EGFR-Viii BaF3 transformants was similar to that for EGFR-E746-750 mutant cells (39nM versus 41nM), albeit much less potent versus the IC50 observed for tumor cells driven by HER2-WT (IC50 1.6nM for BT474 tumor cells).

Since covalent dimerization of LoDi-EGFR oncogenes was potentiated by small molecules that bound to the active conformation of receptor, we speculated that a catalytic domain mutation which also stabilizes the active conformation of the kinase domain would similarly promote covalent dimerization of LoDi-EGFR. We engineered the EGFR-E746-750 active site mutation in cis with EGFR-Viii, as this deletion mutation has been shown to activate the kinase by conferring the active conformation (27). Just as the Type I inhibitor erlotinib promotes covalent dimerization of EGFR-Viii, the presence of the E746-750 mutation, in cis with EGFR-Viii, also promotes covalent dimerization (Figure 5E). These studies further demonstrate how “inside-out” signaling mediated by either mutations or small molecules that confer the active kinase conformation stimulate covalent dimerization within the extracellular domain of LoDi-EGFR oncogenes.

### LoDi-EGFR oncogenes are activated as covalent dimers in patient derived xenografts, exhibiting altered pharmacology for Type I and II inhibitors

Next, we validated the covalent dimer activation mechanism and receptor pharmacology for LoDi-EGFR oncogenes expressed in patient derived xenografts (PDX). GBM6 is a glioblastoma tumor previously shown to express the EGFR-Viii mutation (28), which we confirmed by RT-PCR (data not shown). In GBM6 PDX tumors we found the presence of a covalently activated EGFR-Viii when tumor lysates were resolved by SDS-PAGE under non-reducing conditions, and this could be reduced to the monomeric form in the presence of DTT reductant (Figure 6A). In an acute dose pharmacodynamics study, while neratinib (at 50mpk, MTD) effectively inhibited the activation of EGFR-Viii in GBM6 PDX tumors in a sustained manner for greater than 24 hours following acute dosing, erlotinib (at 100mpk, MTD) only transiently inhibited EGFR-Viii activity, which was followed by paradoxical activation of LoDi-EGFR-Viii at 24 hours and 48 hours when drug exposure reached trough levels. When erlotinib was tested at the lower dose of 10mpk, corresponding to the ED50 for EGFR-WT tumors (29), we found that erlotinib was ineffective against inhibition of EGFR-Viii (Figure 6B and Supplementary Figure 3). Therefore, the paradoxical activation by the reversible Type I inhibitor (erlotinib) versus the covalent Type II inhibitor (neratinib), could be demonstrated in PDX tumors similar to in vitro studies in engineered BaF3 tumor cells.

**Figure 6.**
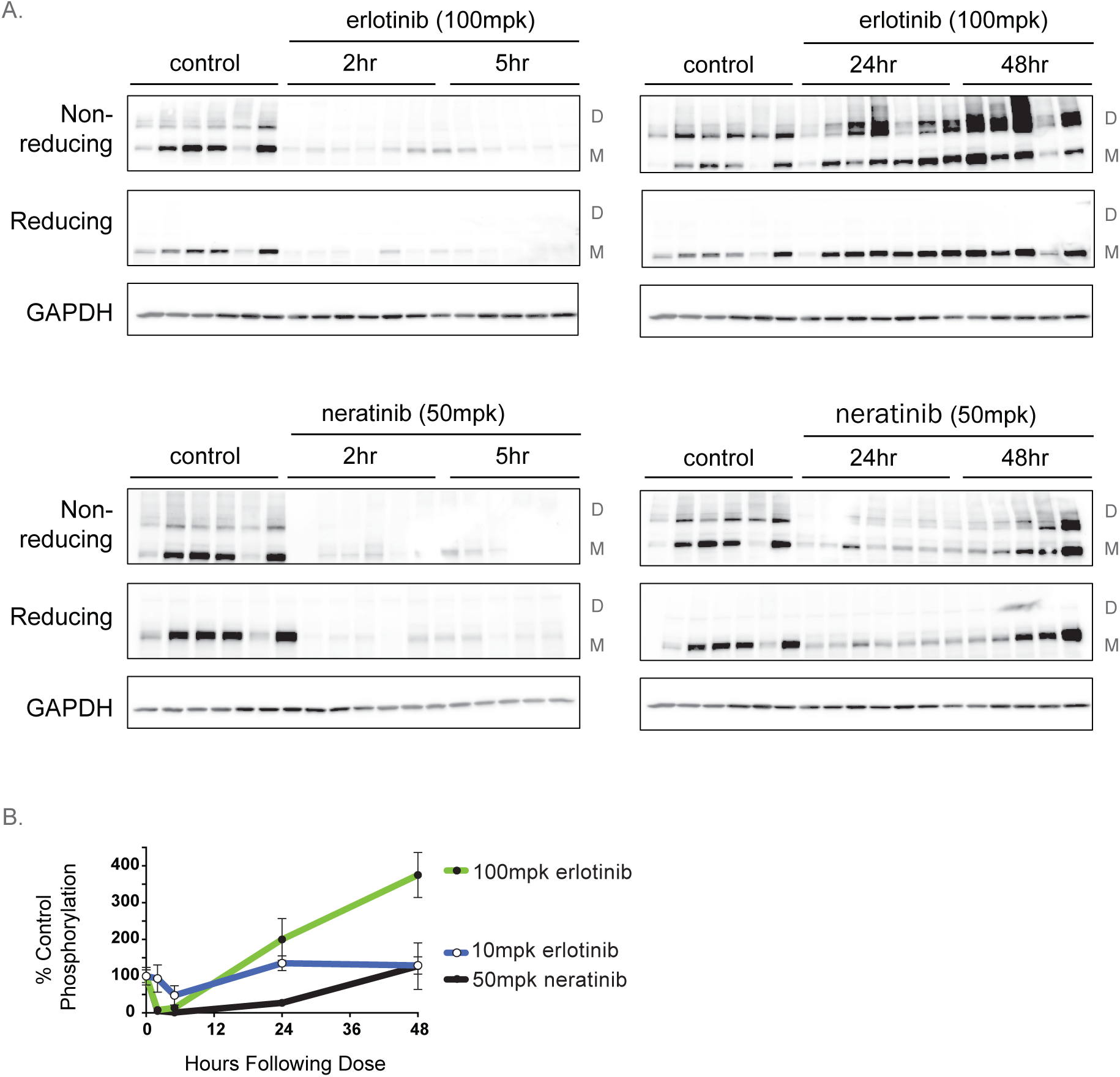
Erlotinib promotes paradoxical activation of EGFR-Viii in PDX tumors. A. Effect of acute dose treatment of erlotinib (100mpk) or neratinib (50mpk) to athymic nude mice bearing GBM6 patient derived tumors expressing EGFR-Viii on levels of total and phosphorylated EGFR. Tumors were collected at varying time points (2, 5, 24, 48 hours) following drug treatment. Tumor lysates were resolved under both non-reducing conditions (to detect covalent dimers) and reducing conditions. B. Densitometric quantitation of immunoblots were performed using AlphaEaseFC software and normalized to GAPDH. Normalized control samples were averaged and considered as maximum signal of 100%.

## Discussion

The extracellular domain of EGFR is an allosteric hot spot for oncogenic mutations in GBM (1,10). Approximately 50% of GBM tumors harbor one or more extracellular domain mutation, and most tumors co-express multiple oncogenic variants. Expression of EGFR mutants tends to be mutually exclusive with expression of other RTK oncogenes, which are co-expressed with EGFR variants in only 7% of GBM tumors (30), demonstrating how EGFR oncogenes in GBM have a dominant and mutually exclusive expression pattern compared with other oncogenic drivers.

Although EGFR mutations in GBM occur at distinct sites of the extracellular domain, spanning a region coded by 10 exons, nearly all mutations affect either of two cysteine rich regions that code for the extracellular dimer interface. We show that a group of the most commonly occurring EGFR truncations in GBM (EGFR-Viii, EGFR-Vii, EGFR-Vvi) all disrupt intramolecular disulfide bonds along this cysteine rich region, resulting in the presentation of free cysteines at the dimerization interface and oncogenic activation by a common mechanism involving disulfide bond mediated covalent homodimerization, or locked-dimerization. We also found evidence for covalent dimerization for EGFR-A289V, the most commonly expressed point mutation expressed in GBM, suggesting that this mutation also disrupts the coupling of intra-molecular disulfide bonds at the dimer interface.

Other studies have attributed the oncogenic activation of EGFR-Viii to the disruption of auto-inhibitory tethering of the extracellular domain, as key interactions at the CR1-CR2 contact point that control the inactive tethered conformation will be lost as a result of this truncation (21). Indeed, auto-inhibitory tethering is also likely disrupted for the EGFR-Vvi oncogene as the tethering contact point at the CR2 site is lost. However, our findings herein indicate that covalent dimerization, and not disruption of auto-inhibition alone, is the primary determinant of oncogenicity. This is supported by data showing the active phosphorylated form of LoDi-EGFR oncogenes is nearly exclusively present as a covalent dimer and how mutating the free cysteine generated by the EGFR-Viii mutation to serine dramatically reduces the activity for EGFR-Viii even though auto-inhibitory tethering would likely be similarly disrupted in both forms. These studies revealed that a diverse collection of allosteric mutations of EGFR in GBM are convergent in both their location at the extracellular dimer interface, and their oncogenic activation through covalent dimerization. This novel mechanism is divergent from the most common EGFR mutations expressed in lung adenocarcinoma, that nearly exclusively affect the intracellular kinase domain. The oncogenic activation of an RTK by distinct groups of mutations has previously been described for other RTKs including RET (rearranged during transfection) (31). As with LoDi-EGFR mutants in GBM, extracellular domain RET mutants in MEN2A patients are predominantly activated by covalent homodimerization, in contrast to patients with MEN2B, where RET mutations virtually exclusively affect the kinase domain. Since cysteine rich domains commonly occur at dimerization interfaces of receptors it is mind-bending to consider how many other RTKs might also be subject to oncogenic mutations that promote covalent homodimerization.

Mutations affecting the extracellular regions of EGFR could be exploited for the design of highly selective biologics or immunotherapeutics. However, since many GBM tumors show a heterogeneic expression pattern for multiple LoDi-EGFR oncogenes, a small molecule that inhibits a collection of variants will likely deliver a greater breadth and duration of treatment benefit compared to an EGFR antibody that is selective only for one isoform. Attempts to reposition small molecule EGFR inhibitors that are approved for the treatment of lung cancer patients expressing EGFR ATP-site mutations have repeatedly failed in GBM patients (14,16). In this study, we demonstrate resistance of LoDi-EGFR oncogenes to these current generation drugs in preclinical models, and reveal the underlying molecular mechanism responsible for their inactivity.

Prior studies have demonstrated that small molecule inhibitors binding at the intracellular ATP site can promote dimerization of EGFR. This induced dimerization occurs at both the asymmetric dimer interface of the intracellular kinase domains and at the symmetric dimer interface of the extracellular domains (32,33). Dimerization induced by small molecules is observed for those that bind to the ATP-site in the active conformation (Type I) with the alphaC helix in the inward state, but not those that bind to the ATP site in the inactive conformation, with outward positioning of the alpha-C helix (Figure 7). We show here that this property for EGFR to undergo “inside-out” signaling has a profound impact on the pharmacology for small molecule inhibitors binding to the ATP-site of LoDi-EGFR oncogenes, even though the primary amino acid sequence of the catalytic site for these mutants is unaffected. Erlotinib, and other small molecules that interact through a Type I binding mode, potentiate covalent dimerization for LoDi-EGFR oncogenes, an observation made for all 4 LoDi-EGFR oncogenes tested and validated in patient derived xenografts expressing EGFR-Viii. Dimer induction is unique to Type I molecules that bind to the active conformation of the kinase domain, and not observed with Type II molecules that bind to the inactive conformation. Here, dimer induction is related to the ability of select small molecules to stabilize the “alpha-C helix in” active conformation, as covalent dimerization is similarly induced when mutations conferring the active alpha-C helix in conformation are engineered in cis with EGFR-Viii.

**Figure 7.**
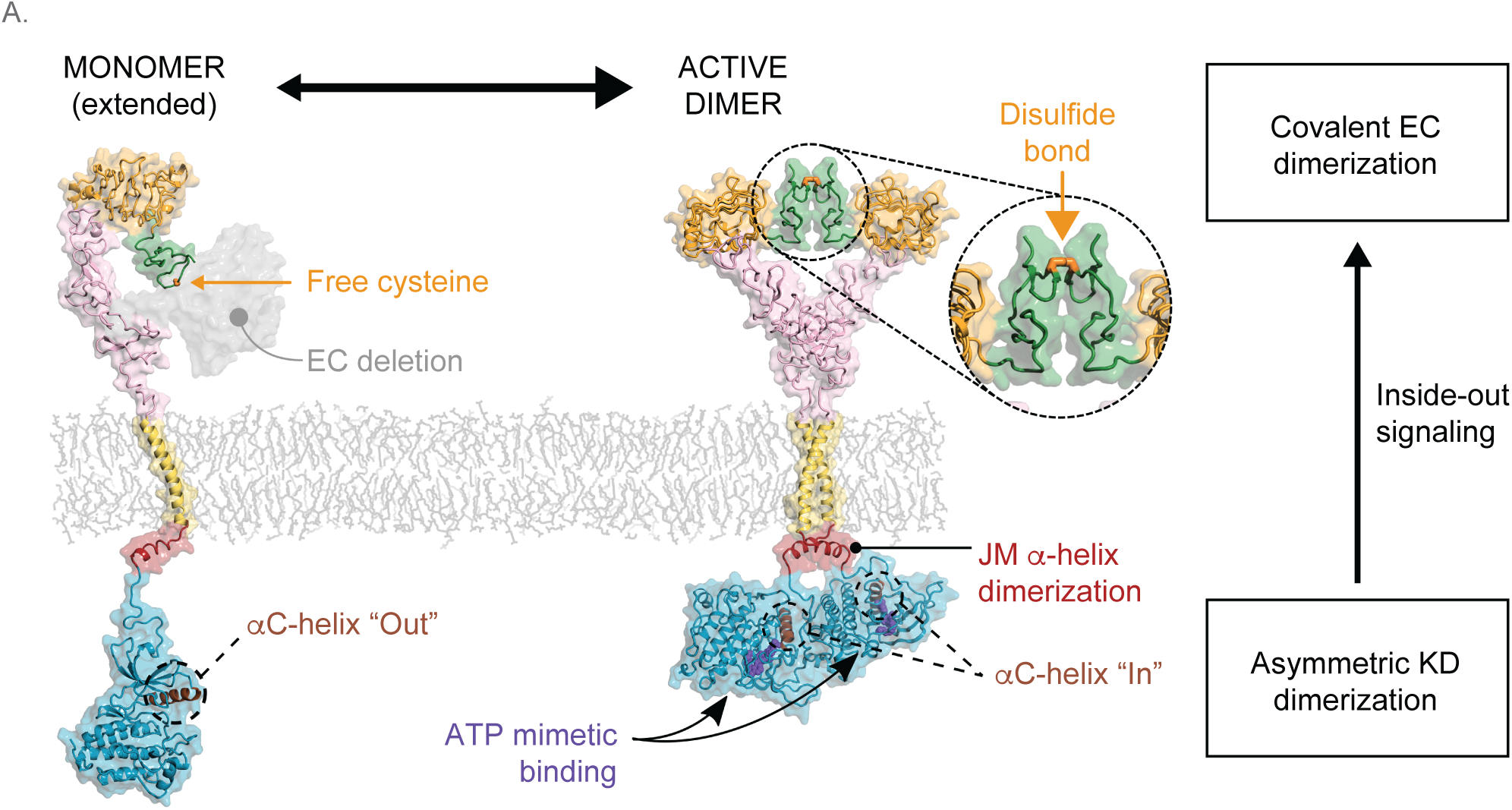
ATP mimetics that bind to the active conformation of EGFR promote covalent dimerization of LoDi-EGFR oncogenes because of “inside-out” EGFR signaling. EGFR is inactive as a monomer (left panel). Oncogenic mutations that result in free cysteines at the extracellular (EC) domain dimer interface lead to ligand-independent activation of a covalently dimerized receptor. Small molecule ATP mimetics that bind to the intracellular kinase domains (KD) with the active αC-helix “in” conformation stabilize asymmetric dimerization, a conformation that leads to enhanced covalent dimerization of the extracellular domain for LoDi-EGFR mutants.

Dimer induction by Type I inhibitors is associated with paradoxical activation of receptor activity and proliferation for LoDi-EGFR transformants. Erlotinib treatment results in a “bell-shaped” dose response profile for each LoDi-EGFR mutant cell line, wherein sub-saturating concentrations of inhibitor stimulate cell proliferation. Paradoxical activation for erlotinib was similarly seen in the EGFR-Viii PDX tumor model, where in an acute dose study, treatment with erlotinib resulted in activation of EGFR-Viii phosphorylation when drug levels reached trough levels. The paradoxical activation of kinase activity by small molecules in response to dimer induction has similarly been seen for BRAF (34). In this case, the binding of drugs such as vemurafenib to the ATP site results in induced dimerization of BRAF under cellular conditions that provide a supportive environment for dimerization, i.e. elevated Ras signaling. In turn, this induced dimerization results in paradoxical activation of BRAF at sub-saturating doses of inhibitor, the so called “BRAF paradox” (34). Our observations for paradoxical activation of EGFR shares parallels with the BRAF paradox, and is the first time to our knowledge of this effect being observed for an RTK.

In contrast to Type I inhibitors, we found that Type II inhibitors, including neratinib, are devoid of paradoxical activation for cells expressing LoDi-EGFR oncogenes, which is consistent with the absence of induced dimerization for this inhibitor class. We captured these observations in PDX tumors expressing LoDi-EGFR-Viii where neratinib inhibited EGFR phosphorylation at all time points tested, albeit this inhibition was observed when neratinib was dosed at a level producing nearly ten times the exposure attained in patients at the maximum tolerated dose. Neratinib’s potency against LoDi-EGFR mutants is at least 20-fold weaker versus HER2-WT, the receptor associated with neratinib’s clinical activity, and on par with EGFR-E746-750, a mutant that has been associated with non-responsiveness to neratinib in patients. As such, to simply reposition neratinib to treat the LoDi-EGFR patient population is not advisable.

Our innovative findings provide a mechanistic understanding for how structural variations affecting EGFR regions distal to the active site can confer dramatically different responses to small molecule active site inhibitors. Our findings for paradoxical activation of covalently-activated EGFR oncogenes by Type I inhibitors have several important clinical implications. Firstly, our findings finally provide a mechanistic explanation for the failed clinical studies for Type I inhibitors, including erlotinib and gefitinib, in GBM. Secondly, pre-screening of patients for tumor expression levels for LoDi-EGFR mutants before treatment with Type I ErbB inhibitors may be warranted, and may even become an exclusion criteria for this type of therapy. Finally, this new understanding may prove to be a catalyst for the optimization of selective Type II inhibitors that are targeted against LoDi-EGFR oncogenes in GBM.

## Methods

### Retroviral Production

EGFR mutants were subcloned into pMXs-IRES-Blasticidin (RTV-016, Cell Biolabs, San Diego, CA). Retroviral expression vector retrovirus was produced by transient transfection of HEK 293Tcells with the retroviral EGFR mutant expression vector pMXs-IRES-Blasticidin (RTV-016, Cell Biolabs), pCMV-Gag-Pol vector and pCMV-VSV-G-Envelope vector. Briefly, HEK 293T/17 cells were plated in 100mm plates (430293, Corning, Tewksbury, MA) (4 x 10^5^ per plate) and incubated overnight. The next day, retroviral plasmids (3µg of EGFR mutant, 1.0 µg of pCMV-Gag-Pol and 0.5µg pCMV-VSV-G) were mixed in 500 µl of Optimem (31985, Life Technologies). The mixture was incubated at room temperature for 5 min and then added to Optimem containing transfection reagent Lipofectamine (11668, Invitrogen) and incubated for 20 minutes. Mixture was then added dropwise to HEK 293T cells. The next day the medium was replaced with fresh culture medium and retrovirus was harvested @ 24 and 48hrs.

### Cell Culture

Ba/F3 cells were purchased from DSMZ (ACC-300, Braunschweig, Germany) and maintained in RPMI1640 medium (11875093Life Technologies) supplemented with 10% FBS, 1% L-glutamine and 10ng/ml mouse IL-3 (203-IL-010, R&D Systems). Ba/F3 transformed cells were maintained in the medium with 15ug/ml Blasticidin S HCl (A1113903, Life technologies). U87MG cells were purchased from ATCC (HTB-14, Manassas, VA) and maintained MEM medium (11095080, Life Technologies) and supplemented with 10%FBS and 1% L-glutamine. U87MG-EGFR mutant overexpressed cells were maintained in the medium with 10ug/ml Blasticidin S HCl (A1113903, Life technologies)

### Generation of EGFR mutant stable cell lines

BaF3 cells (5.0E6 cells) were infected with 2ml of viral supernatant supplemented with 8ug/ml polybrene by centrifuging for 30 min at 1000rpm. Cells were placed in a 37°C incubator overnight. Cells were then spun for 5 minutes to pellet the cells. Supernatant was removed and cells re-infected a fresh 2ml of viral supernatant supplemented with 8ug/ml polybrene by centrifuging for 30min at 1000rpm. Cells were placed in 37°C incubator for 4hrs. Cells were then maintained in RPMI containing 10% Heat Inactivated FBS, 2% L-glutamine containing 10ng/ml IL-3. After 24hrs cells were selected for retroviral infection in 10ug/ml Blasticidin for one week. Blasticidin resistant populations were washed three times in phosphate buffered saline before plating in media lacking IL-3 to select for IL-3 independent growth.

### Measurement of cell proliferation

BaF3 cell lines were resuspended at 1.1E5 c/ml in RPMI containing 10% Heat Inactivated FBS and 2% L-glutamine and dispensed in triplicate (1.485E4 c/well) into 96 well plates. To determine the effect of drug on cell proliferation, cells incubated for 3 days in the presence of vehicle control or test drug at varying concentrations. Inhibition of cell growth was determined by luminescent quantification of intracellular ATP content using CellTiterGlo (Promega), according to the protocol provided by the manufacturer. Comparison of cell number on day 0 versus 72 hours post drug treatment was used to plot dose-response curves. The number of viable cells was determined and normalized to vehicle-treated controls. Inhibition of proliferation, relative to vehicle-treated controls was expressed as a fraction of 1 and graphed using PRISM^®^ software (Graphpad Software, San Diego, CA). EC_50_ values were determined with the same application

### Cellular protein analysis

Cell extracts were prepared by detergent lysis (RIPA, R0278, Sigma, St Louis, MO) containing 10mM Iodoacetamide (786-228, G-Biosciences, St, Louis, MO), protease inhibitor (P8340, Sigma, St. Louis, MO) and phosphatase inhibitors (P5726, P0044, Sigma, St. Louis, MO) cocktails. The soluble protein concentration was determined by micro-BSA assay (Pierce, Rockford IL). Protein immunodetection was performed by electrophoretic transfer of SDS-PAGE separated proteins to nitrocellulose, incubation with antibody, and chemiluminescent second step detection. Nitrocellulose membranes were blocked with 5% nonfat dry milk in TBS and incubated overnight with primary antibody in 5% bovine serum albumin. The following primary antibodies from Cell Signaling Technology were used at 1:1000 dilution: phospho-EGFR[Y1173] and total EGFR. β-Actin antibody, used as a control for protein loading, was purchased from Sigma Chemicals. Horseradish peroxidase-conjugated secondary antibodies were obtained from Cell Signaling Technology and used at 1:5000 dilution. Horseradish peroxidase-conjugated secondary antibodies were incubated in nonfat dry milk for 1 hour. SuperSignal chemiluminescent reagent (Pierce Biotechnology) was used according to the manufacturer’s directions and blots were imaged using the Alpha Innotech image analyzer and AlphaEaseFC software (Alpha Innotech, San Leandro CA).

### Analysis of solubilized and purified EGFR variants

FreeStyle 293-F cells were purchased from Life Technologies and maintained in FreeStyle 293 expression medium (Life technologies, 12338026). EGFR variants C-terminally fused to EGFP, followed by His10-tag and Twin-Strep-tag were cloned into a mammalian expression vector, under the control of CMV promotor. In the expression constructs for large scale expression, EGFP was omitted and the EGFR genes contained only C-terminal His10-tag and Twin-Strep tag. The expression plasmids were transiently transfected into 293-F cells (at the density of 1E6 c/ml) using polyethyleneimine (Polysciences Inc., 23966-1) and Opti-MEM I reduced serum medium (Life Technologies, 1058021). For the analysis of EGFR-EGFP fusions, 2 ml suspension cultures were grown in 24-well plates. When erlotinib effect was tested, this inhibitor was added to the wells 5 hours after transfection at 2, 4, 6, 8 and 10 μM final concentration. 48 h after transfection the cells were harvested by centrifugation, and washed with PBS containing 10 mM iodoacetamide. Cell pellets (10^6^ cells) were solubilized for 1hour at 4°C in 20 mM Tris-HCl pH 8, 200 mM NaCl, 1% DDM, 2 mM iodoacetamide supplemented with protease inhibitors. Solubilized cell lysates were cleared by centrifugation and supernatants analyzed by fluorescence-detection size exclusion chromatography on a Superdex 200 Increase 5/150 column equilibrated in 20 mM Tris-HCl pH 8, 200 mM NaCl, 0.025% DDM. EGFP fluorescence (excitation 488 nm, emission 510 nm) of eluted samples was monitored using Agilent 1260 Infinity II fluorescence detector. For obtaining in-gel fluorescence data, the same samples were mixed with non-reducing sample buffer, loaded onto SDS-PAGE and visualized for fluorescence using a gel imager.

For purification of solubilized EGFRvIII, 1 L culture of transfected FreeStyle 293-F cells was used. Erlotinib concentration in large scale expression was 5 μM. After harvesting, cells were solubilized for 1hour at 4°C in the buffer: 50 mM Tris-HCl pH 8, 200 mM NaCl, 1 mM EDTA, 1% DDM, 2 mM iodoacetamide supplemented with protease inhibitors. The suspension was centrifuged using Beckman Coulter Optima XP-100 ultracentrifuge for 45 min at 60’000 rpm, in the rotor Ti70 at 4°C. Solubilized EGFRvIII was purified from the supernatant using affinity chromatography on Strep-Tactin XT resing (IBA-LifeSciences) using procedure provided by the manufacturer. Eluted protein was concentrated using VIVASPIN TURBO 15 concentrator with 100 kD MWCO and injected onto Superdex 200 10/300 Increase column equilibrated in 20 mM Tris-HCl pH 8.0, 200 mM NaCl, 1 mM EDTA, 0.025% DDM. The eluted fractions were analyzed by SDS-PAGE.

### Determination of subcutaneous growth of EGFR mutant cell lines in athymic nude mice

Five-week-old female athymic nude mice (Charles River Laboratories nude mice (CRL:NU-Foxn1nu)) were implanted with 5X10^6 EGFR-Viii or EGFR-Ex10del BaF3 transformant cells in 50% matrigel subcutaneously on the high right axilla. Tumor volume was measured 3X per week.

### PDX tumor studies

Five-week-old female athymic mice (nu/nu, BALB/c background) purchased from Simonsen Laboratories (Gilroy, CA) were used. Animals were housed under aseptic conditions. All animal protocols were approved by the UCSF Institutional Animal Care and Use Committee. GBM6 patient derived xenografts were purchased from the Brain Tumor Center Preclinical Therapeutic Testing Core in UCSF. Tumor chunks were subcutaneously implanted into both left and right flanks of athymic mice. These mice were randomized into four groups receiving treatment by oral gavage: (1) vehicle control (6% Captisol, and 0.5% hydroxypropyl methylcellulose plus 0.2% Tween 80); (2) 10 mg/kg Erlotinib (dissolved in 6% Captisol); (3) 100 mg/kg Erlotinib (dissolved in 6% Captisol); (4) 50 mg/kg neratinib (dissolved in 0.5% hydroxypropyl methylcellulose plus 0.2% Tween 80). Mice in the vehicle control group were sacrificed 2 hours after treatment on the second day, and mice in other groups were sacrificed at the time points of 2, 5, 24 and 48 hours after treatment on the second day (3 mice for each time point). Tumors, brain, and plasma of each mouse were collected and snap frozen in liquid nitrogen for subsequent analysis. All treatments were initiated when tumors reach a size of 300 mm^3^. The maximum longitudinal diameter (length) and the maximum transverse diameter (width) of tumors were measured by caliper every day, and tumor volume was calculated using the equation 1/2 × (length × width^2^).

