## Supplemental Figures for "EGFR Oncogenes Expressed In Glioblastoma Are Activated As Covalent Dimers And Paradoxically Stimulated By Erlotinib"

**Supplementary Figure 1. Type I, but not Type II, ErbB inhibitors enhance the formation of covalent dimers for the LoDi-EGFR mutant EGFR-A289V**

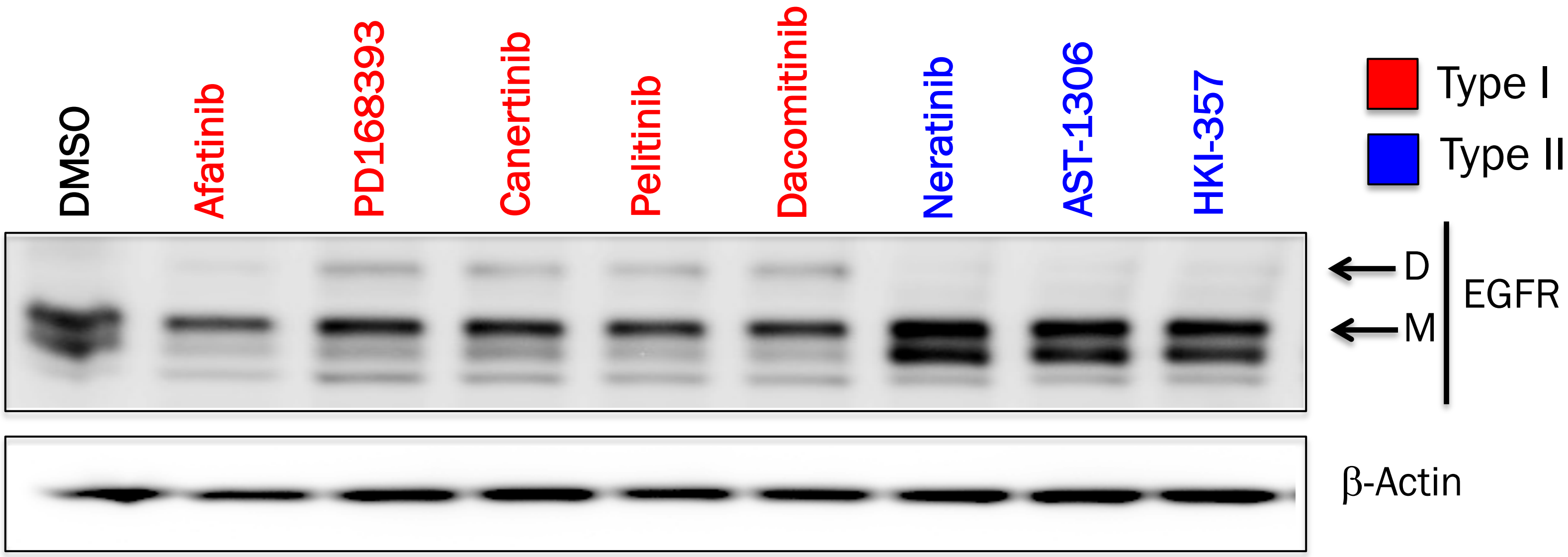

**Supplementary Figure 2. Gefitinib stimulates the proliferation of cells driven by EGFR-Viii**

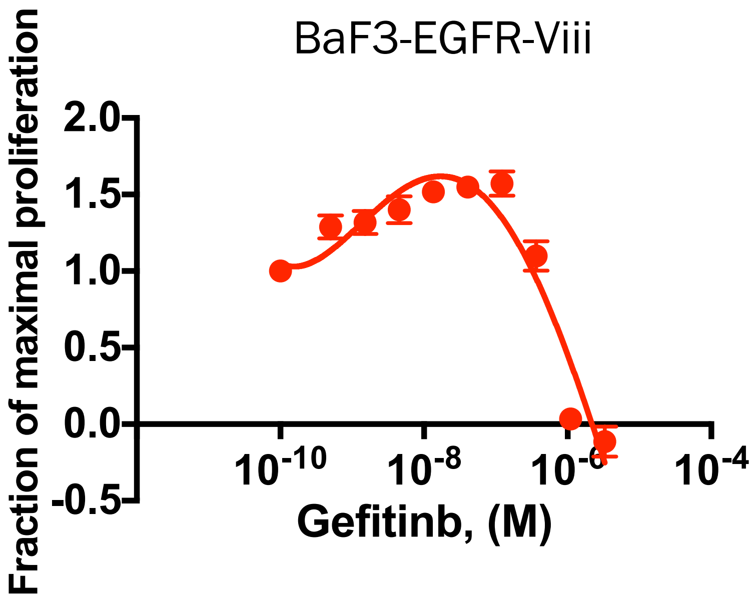

### Supplementary Figure 3.

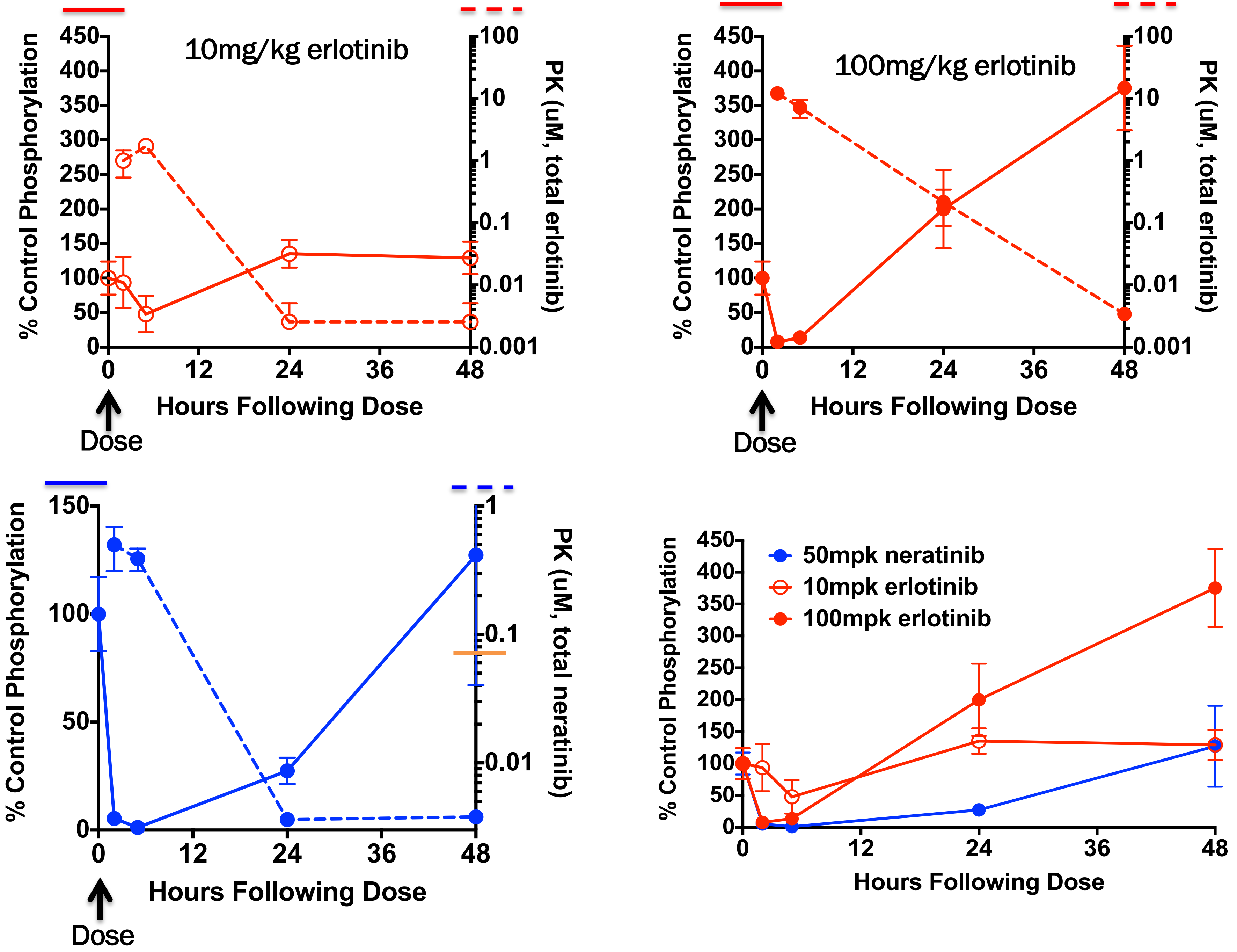
