## Supplementary Figure Legends for "EGFR Oncogenes Expressed In Glioblastoma Are Activated As Covalent Dimers And Paradoxically Stimulated By Erlotinib"

**Supplementary Figure 1. Type I, but not Type II, ErbB inhibitors enhance formation of covalent dimers for LoDi-EGFR mutants.** A. Effect of a group of small molecule ErbB inhibitors (100nM) on levels of monomeric and covalently dimerized EGFR-A289V.

**Supplementary Figure 2. Gefitinib potentiates the proliferation of EGFR-Viii BaF3 transformants.** Effect of varying concentrations of erlotinib on the proliferation of BaF3 transformant cells.

**Supplementary Figure 3. Erlotinib, but not neratinib, treatment results in paradoxical stimulation of EGFR-Viii phosphorylation at drug trough levels.** Effect of acute dosing of erlotinib at 10mpk (A) or 100mpk (B) or neratinib at 50mpk (C) on the level of EGFR-Viii phosphorylation in GBM6 tumors. Phosphorylation of EGFR was determined by Western Blotting, and results from quantitative densitometry are shown as % of control phosphorylation in untreated tumors. Also shown is total plasma levels of erlotinib or neratinib at indicated time points. Orange bar on right y-axis in part C indicates the total plasma exposure for neratinib at steady state (Cmax) observed in patients treated with the FDA approved dose of 240mg daily, reported as 73ng/ml in Wong et al (36) and 45.5ng/ml in Nerlynx FDA briefing documents. D. Overlay of phosphorylated EGFR signals shown in parts A-C.
